# Mechanical effects of MitraClip on leaflet stress and myocardial strain in functional mitral regurgitation – A finite element modeling study

**DOI:** 10.1101/687004

**Authors:** Yue Zhang, Vicky Y Wang, Ashley E Morgan, Jiwon Kim, Mark D Handschumacher, Chaya S Moskowitz, Robert A Levine, Liang Ge, Julius M Guccione, Jonathan W Weinsaft, Mark B Ratcliffe

## Abstract

**Purpose:** MitraClip is the sole percutaneous device approved for functional mitral regurgitation (MR; FMR) but MR recurs in over one third of patients. As device-induced mechanical effects are a potential cause for MR recurrence, we tested the hypothesis that MitraClip increases leaflet stress and procedure-related strain in sub-valvular left ventricular (LV) myocardium in FMR associated with coronary disease (FMR-CAD).

**Methods:** Simulations were performed using finite element models of the LV + mitral valve based on MRI of 5 sheep with FMR-CAD. Models were modified to have a 20% increase in LV volume (↑LV_VOLUME) and MitraClip was simulated with contracting beam elements (virtual sutures) placed between nodes in the center edge of the anterior (AL) and posterior (PL) mitral leaflets. Effects of MitraClip on leaflet stress in the peri-MitraClip region of AL and PL, septo-lateral annular diameter (SLAD), and procedure-related radial strain (*E*_*rr*_) in the sub-valvular myocardium were calculated.

**Results:** MitraClip increased peri-MitraClip leaflet stress at end-diastole (ED) by 22.3±7.1 kPa (p<0.0001) in AL and 14.8±1.2 kPa (p<0.0001) in PL. MitraClip decreased SLAD by 6.1±2.2 mm (p<0.0001) and increased *E*_*rr*_ in the sub-valvular lateral LV myocardium at ED by 0.09±0.04 (p<0.0001)). Furthermore, MitraClip in ↑LV_VOLUME was associated with persistent effects at ED but also at end-systole where peri-MitraClip leaflet stress was increased in AL by 31.9±14.4 kPa (p=0.0268) and in PL by 22.5±23.7 kPa (p=0.0101).

**Conclusions:** MitraClip for FMR-CAD increases mitral leaflet stress and radial strain in LV sub-valvular myocardium. Mechanical effects of MitraClip are augmented by LV enlargement.

## Introduction

Functional mitral regurgitation (FMR) in patients with coronary artery disease (FMR-CAD) occurs in 1.2 to 2.1 million patients in the United States and of those more than 400,000 patients have advanced (≥2+) FMR-CAD. [1] These numbers are expected to progressively increase as the population ages and more patients survive acute myocardial infarction (MI). [1] Advanced FMR-CAD discovered at cardiac catheterization has a 1-year mortality of approximately 17%. [2] One-year mortality for severe (≥3+) FMR-CAD is approximately 40%. [2]

MitraClip (Abbott Vascular, Santa Clara, CA) is the sole percutaneous device commercially approved for treatment of FMR. Over 80,000 patients have undergone mitral repair with the MitraClip device in the past decade. [7]. The device approximates the anterior (AL) and posterior leaflets (PL) of the mitral valve, creating a double-barrel mitral orifice as a means of decreasing mitral regurgitation. [3] Two recent trials (COAPT, MITRA-FR) of MitraClip in patients with FMR reported discordant results. MITRA-FR found no difference in mortality and congestive heart failure (CHF) [4], whereas COAPT found that mortality and CHF were lower in MitraClip treated patients. [5]

A key issue with MitraClip is recurrence of MR where recurrent advanced MR is present in over one third of FMR patients treated with MitraClip. [8-10] Mechanisms responsible for MR recurrence after MitraClip for FMR remain unclear. However, there is reason to believe that LV chamber volume is a primary determinant of recurrent MR after MitraClip. For instance, LV chamber volume was over 25% greater among patients in MITRA-FR (135 ± 35 ml/m^2^) as compared to COAPT (101 ± 34 ml/m^2^), paralleling increased rates of residual MR. [4, 5] These studies suggest a link between LV chamber size and outcome mediated by mechanical effects of MitraClip including mitral leaflet stress and procedure related tissue stretch in sub-valvular LV myocardium.

Finite element (FE) based computational modeling allows calculation of mitral leaflet stress. For instance, Votta et al. used an FE model of the mitral valve to simulate the effect of edge-edge repair and showed that leaflet stress increases with annular dilatation. [12] Sturla et al. simulated MitraClip in an FE model of human PL prolapse and reported that leaflet stress at peak systole after MitraClip is dependent on LV pressure and that stress was increased with asymmetric device application. [13] Recently, Morgan et al. simulated a FE model of human posterior leaflet prolapse that included the LV + mitral valve and reported that uneven grasp of leaflet tissue by the MitraClip did not increase leaflet stress [14], as well as stresses and strains in the mitral apparatus and adjacent LV wall. To date, however, there have been no FE models of MitraClip in FMR.

This study employed FE models of the LV + mitral valve based on sheep with FMR after postero-lateral MI [15, 16] to test the hypothesis that MitraClip increases mitral leaflet stress and procedure-related strain in the sub-valvular LV myocardium..

## Methods

Animals used for creating the finite element models were treated in compliance with the “Guide for the Care and Use of Laboratory Animals”. [17] Adult sheep underwent postero-lateral MI as previously described. [18] Cardiac magnetic resonance imaging (MRI) showed that infarct area was 21 ± 7 % [16] and animals developed mild to moderate FMR-CAD 16 weeks after MI. [18]

### Model construction and constitutive equations

The LV was contoured and meshed as previously described. [19] Rule-based fiber angles were assigned with myofiber fiber helix angle varying transmurally from −60° at the epicardium to +60° at the endocardium. LV myocardium was divided into MI, borderzone, and remote regions (**Figure 1A**). Active and passive myocardial constitutive laws were previously described by Guccione et al. [20, 21] Prior to virtual MitraClip, the passive stiffness parameter, *C*, and contractility parameter, *T*_*max*_, for MI, borderzone and remote regions were inversely calculated by minimizing the difference between modeled and experimentally determined LV volume and regional strain [22] using established methods. [23] The optimized values for *C* and *T*_*max*_ for MI, borderzone and remote regions were previously reported. [15]

**Figure 1.**
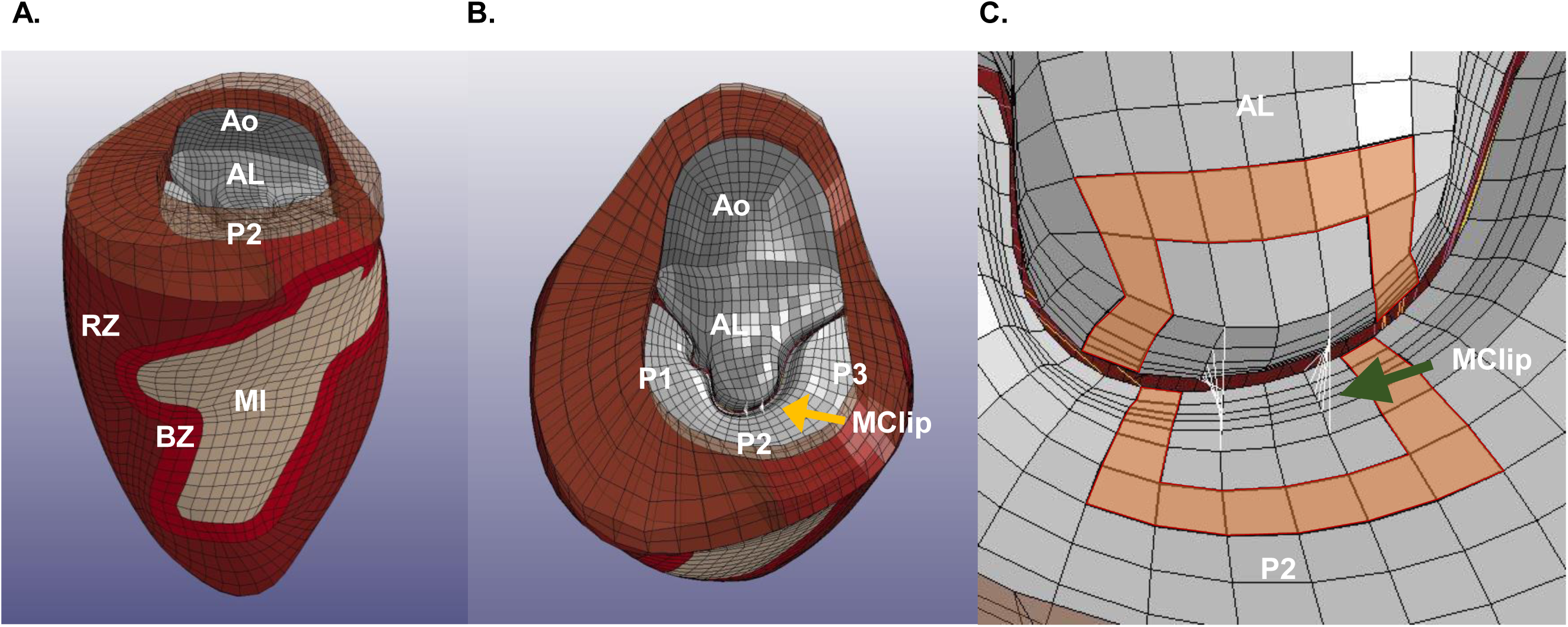
Representative FE mesh of the LV with Mitral Valve and MitraClip modeled as virtual sutures. Views include (**A**) LV showing postero-lateral MI, (**B**) en face view of the mitral valve with MitraClip and (**C**) close up of MitraClip as virtual sutures. The leaflet elements used for stress calculation are marked in orange. BZ – infarct borderzone; RZ – remote myocardium; AL – anterior leaflet; P1,2,3 – posterior leaflet sections; Ao – capped region of aortic outflow and MClp – MitraClip.

The mitral leaflets were contoured and meshed as previously described (**Figure 1B**). [19] Edge chords were attached to free edge of each leaflet and strut chords were attached to the mid-section of each leaflet, as described by Wenk et al.. [19] The mitral leaflets were modeled using a bi-layer soft tissue material (*MAT_091, LS-DYNA, Livermore Software Technology Corporation (LSTC), Livermore, CA) [24] and the chordae tendineae were modeled with cable elements (*MAT_071; LSTC).

### Loading and boundary conditions

The endocardial surface of the FE model was loaded with animal-specific in-vivo LV end-diastolic and end-systolic pressures. During diastole, the mitral leaflets were loaded with MVPG of −4.1 mm Hg in line with MVPG data after MitraClip from Herrmann et al. [25] During systole, MVPG equaled measured LV end-systolic (ES) pressure.

To isolate the mitral apparatus from boundary condition restrictions, the myocardial elements at the level of the valve were extended 3 elements above the mitral plane. The extended base had diastolic stiffness, *C*, equal to 0.001\**C* of remote myocardium and *T*_*max*_ equal to zero. Homogeneous Dirichlet boundary conditions were applied to nodes at the top of the extended base so that the nodes were fixed in the longitudinal axis but allowed to slide in the mitral plane.

### Virtual MitraClip

The MitraClip (NTR, Abbott, Abbott Park, Il) was simulated by attaching leaflet nodes in a rectangular pattern extending from the centers of the leading edge of the central regions of the anterior and posterior leaflets (A2 and P2; **Figure 1C**). The average body surface area (BSA) of the 5 sheep on which the simulations were based was 1.28±0.04. Given that the normal human BSA is 1.99 [26] the MitraClip arm dimensions (9 mm height and 5 mm width) were indexed to 5.7 × 3.2 mm. A 70% partial grasp MitraClip device where dimensions were 4.0 × 2.9 mm was also simulated.

Leaflet nodes were attached using ‘virtual suture’ beam elements (material model: *MAT_071, LS-DYNA) as previously described. [27] Specifically, each virtual suture was modeled as a discrete beam element. A unique feature of *MAT_071 is that an axial tension can be specified for each beam element and this axial tension could mechanically pull the two ends of each element together. After the leaflet coaptation, the virtual sutures were changed to rigid elements using the *DEFORMABLE_ TO_RIGID function in LS-DYNA. Virtual MitraClip was performed at the start of diastole (**Figure 2**). Simulation of LV diastole and systole then proceeded.

**Figure 2.**
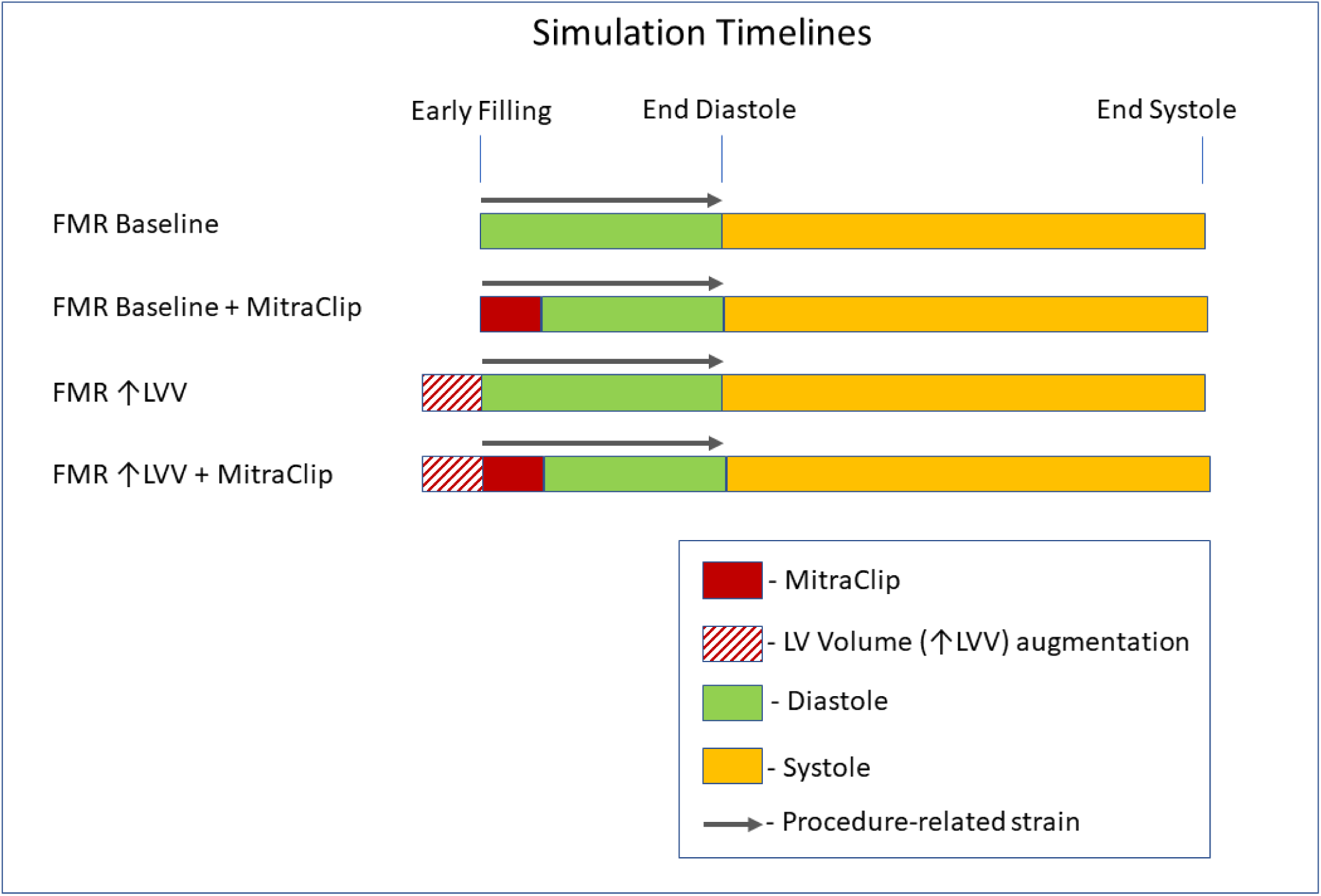
Simulation time lines.

### Simulation of LV volume effect

LV volume at early filling was increased by using a pre-inflation step where the LV endocardial surface was loaded with animal-specific LV ED pressure until an increase in LV volume of 20% was achieved. The deformed FE mesh was used as the unloaded geometry of a new ↑LV_VOLUME model. Simulation of virtual MitraClip was then performed using the new ↑LV_VOLUME model (**Figure 2**).

### Measured outcomes

Von Mises stress was averaged across anterior and posterior leaflets. Von Mises stress was measured in the one element wide region of elements one element removed from the MitraClip (Peri-MitraClip; **Figure 1C**). Elements with nodes attached to virtual suture beam elements were excluded to avoid stress concentrations associated with local MitraClip-induced element distortion.

Green-Lagrange strain was calculated where the reference geometry for strain at ED is the pre-operative early-diastolic filling LV shape. [16] *E*_*rr*_ at ES is relative to ED. Strain was averaged in a transmural group of elements bounded by the P2 scallop of the mitral valve and extending to a point 1/3 of the distance from the valve to apex (lateral sub-valvular).

### Software and hardware

FE **s**imulations were solved with LS-DYNA (R9.2.0, LSTC) running on a standalone Windows (version 10, Microsoft, Redmond, WA) workstation (i9-18 core, 2.60 GHz, Alienware, Dell, Round Rock, TX). Simulations used between 3 and 16 virtual processors depending on resource availability.

### Statistical analysis

All values are expressed as mean ± standard deviation and compared by repeated measures analysis using a mixed model to test for both fixed and random effects (PROC MIXED, SAS Studio, SAS Release 9.04, University Edition, SAS Institute, Cary, NC). Models included categorical factors for MitraClip clip size, ↑LV_VOLUME and leaflet region. ED and ES analyses were done separately and in all simulations the animal/ model number was treated as a random variable. Significance was set at p<0.05.

## Results

All FE simulations finished successfully with an average calculation time of 64.7 ± 24.6 minutes. Linear regression of simulation calculation time (CalcTime) with the number of virtual processors (NCPU) was CalcTime = −4.95 NCPU + 95 and reached an approximate minimum at 8 virtual processors.

### Mitral leaflet stress

Figure 3 shows representative regional leaflet stress color maps of the mitral valve before and after MitraClip with complete grasp. Changes in leaflet stress are most apparent at LV end-diastole (ED) (**Figure 3B**), during which stress is concentrated in the clip region and increased stress occurs between the device and the mitral annulus. Leaflet stress was also increased in the peri-MitraClip region at LV end-systole (ES) (**Figure 3E**) although the effect is less pronounced. Color maps from the volume augmented (20% ↑LV_VOLUME) FE models (**Figures 3C** and **F**) have a noticeable increase in leaflet stress compared to baseline (non-augmented) models.

**Figure 4** shows the quantitative effects of MitraClip on leaflet stress in the anterior and posterior leaflets at baseline and with LV volume augmentation. As shown, end-diastolic leaflet stress adjacent to the device increased in all animals. At baseline, MitraClip increased peri-MitraClip leaflet stress in the anterior mitral leaflet by 544% and in the posterior leaflet by 131 % (both p<0.0001) (**Figures 4A** and **B**). The effect of MitraClip in ↑LV_VOLUME FE models was similar, as evidenced by mean increases of 207 and 85 % respectively (both p<0.0001). Lesser effects were evident with respect to device-induced leaflet stress in areas remote from the MitraClip: The effect on the leaflet remote from the MitraClip was not significant (NS) with the exception of the anterior leaflet LV volume augmented models in which MitraClip caused a slight, albeit significant, increase in end-diastolic leaflet stress (56 %; p=0.04).

Computational analyses consistently demonstrated MitraClip effects on mitral leaflet stress to be of lesser magnitude during LV end-systole, irrespective of whether measurements were performed in baseline or LV volume augmented models. In baseline models, MitraClip increased peri-device end-systolic leaflet stress by 29.6 % (p=NS) in the anterior leaflet, and by 5.8 % (p=NS) in the posterior leaflet (**Figures 4C and D**). LV volume augmentation resulted in slight increments in device-induced effects on systolic leaflet stress immediately adjacent to the MitraClip. In the ↑LV_VOLUME FE models, MitraClip increased peri-device end-systolic leaflet by 37.4 % in the anterior leaflet (p=0.03), and by 23.3 % (p=0.01) in the posterior leaflet (**Figures 4C** and **D**). Device-induced effects on end-systolic leaflet stress remote from the MitraClip were non-significant for both the anterior and posterior leaflet.

**Figure 3.**
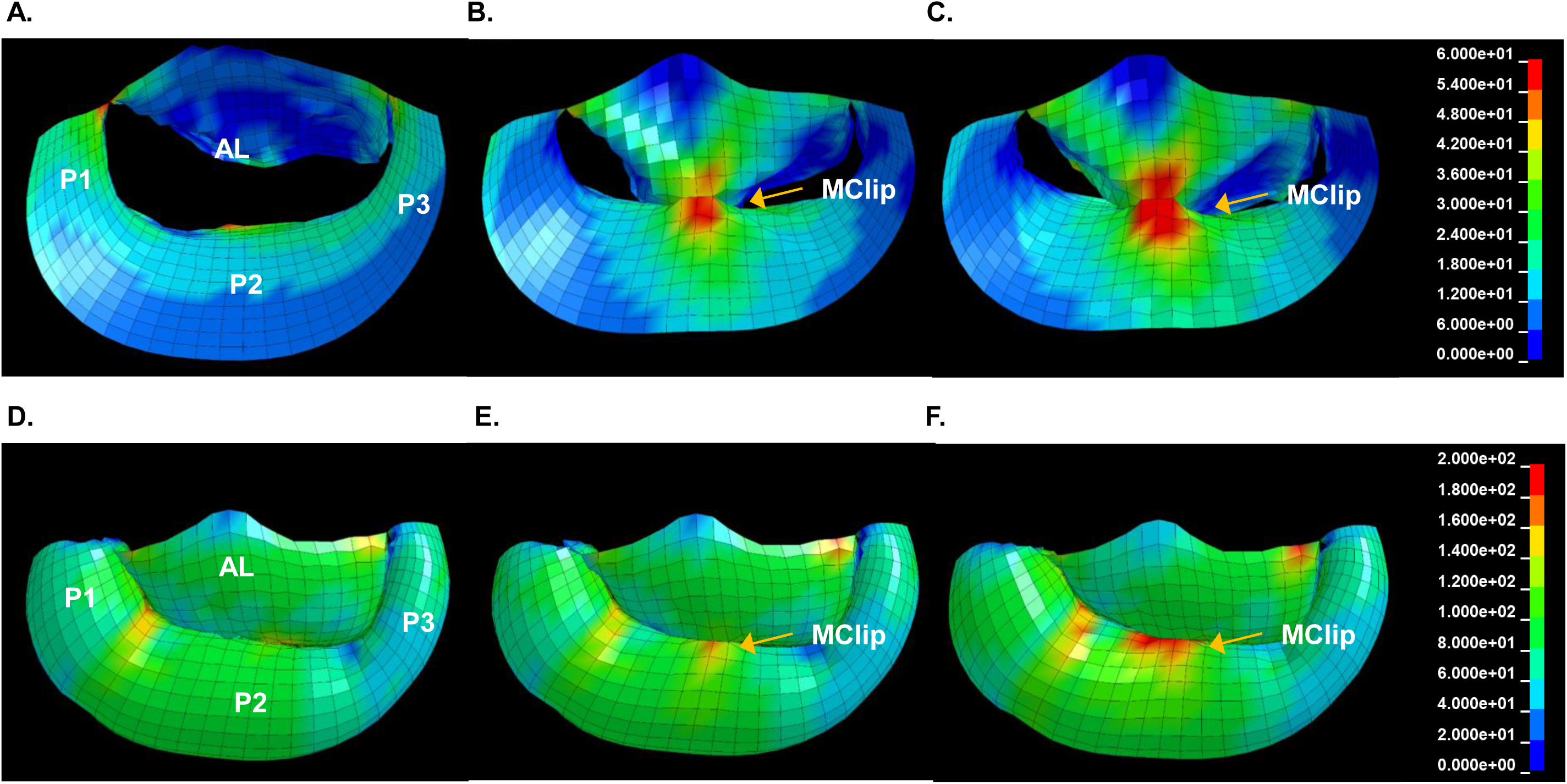
Representative Von Mises stress color maps of the mitral valve before (**A** and **D**) and after (**B** and **E**) virtual MitraClip and after MitraClip in an enlarged LV (**C** and **F**). Effects are seen at end-diastole (**A** - **C**) and end systole (**D** - **F**). Note that these color maps are not dimensionally exact. AL – anterior leaflet; P1,2,3 – posterior leaflet sections; MClip – MitraClip.

**Figure 4.**
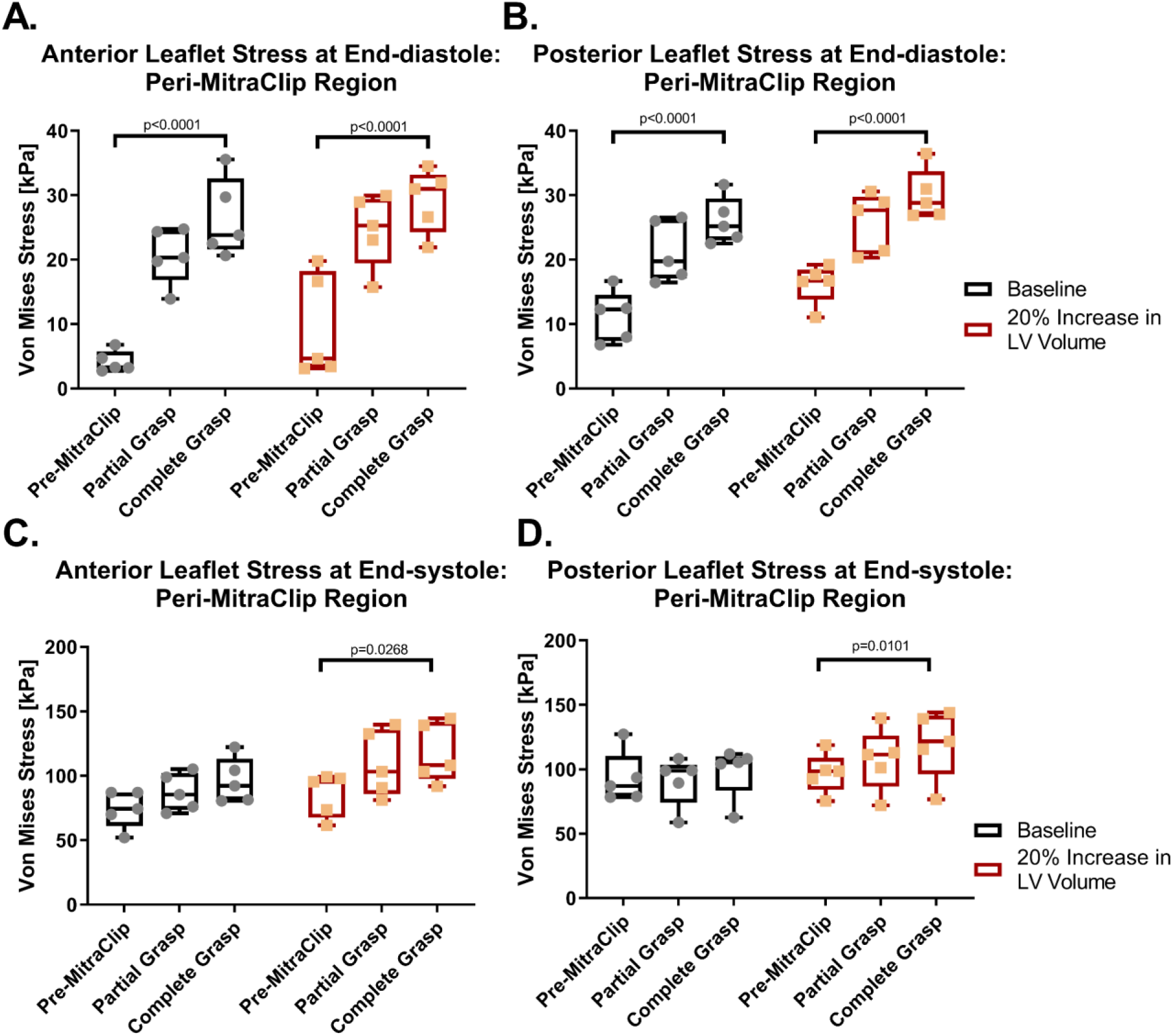
Stress in the peri-MitraClip region in the anterior and posterior leaflets at ED (A and B) and ES (C and D).

Leaflet stress results for the partial grasp technique of MitraClip implantation are also provided in **Figure 4**: As shown, partial grasp impacted leaflet stress in a similar manner to that of complete grasp technique, although device-induced increments in leaflet stress were of lesser magnitude.

### Mitral annular geometry

LV annular diameter, as measured from the septum to the lateral wall (SLAD), before and after MitraClip implantation are shown in **Figure 5**. MitraClip reduced LV end-diastolic SLAD by 18.7 % (p<0.0001) and 21.1 % (p<0.0001) in baseline and LV volume augmented models respectively. Lesser, albeit significant, reductions in LV end-systolic SLAD occurred in both baseline and volume augmented models (8.3 %, and 9.0% % respectively; both p<0.0001). The effect of LV size was not significant at ED but was significant at ES (p=0.03).

**Figure 5.**
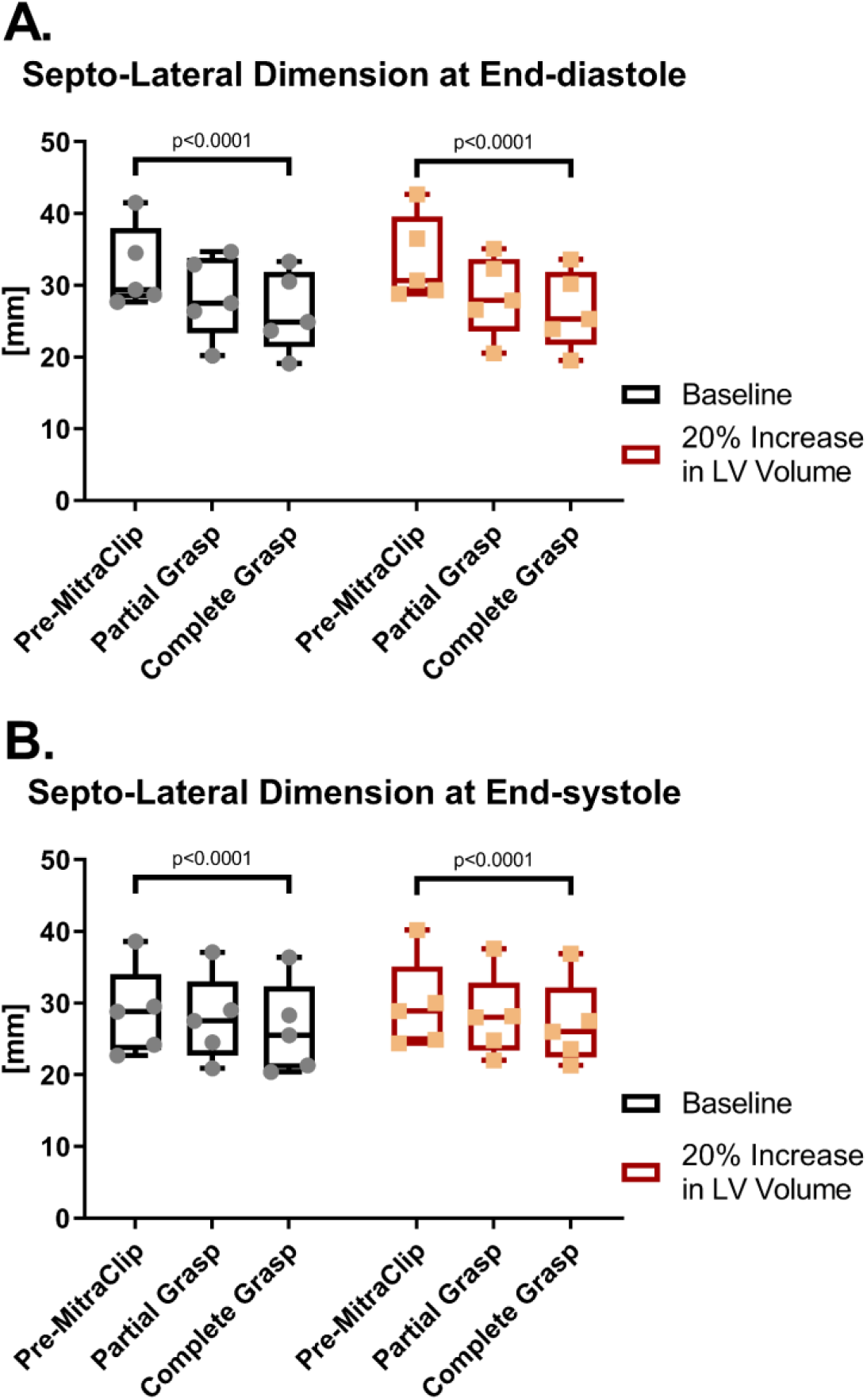
Septo-lateral annular dimension at ED (A) and ES (B) at baseline and after simulated partial and complete grasp MitraClip.

### Sub-valvular myocardial strain

Figure 6 shows representative color maps of change in LV end-diastolic myocardial radial strain (*E*_*rr*_) after simulated MitraClip with complete grasp. Note the change in *E*_*rr*_ in the P2 region (white rectangle) with highest intensity in the P2 endocardium. Note also that intensity is increased in the LV volume augmented model.

Figure 7 illustrates the quantitative effects of MitraClip on radial strain (*E*_*rr*_) in the lateral (P2) sub-valvular myocardium (**Figure 7A**) and endocardium (**Figure 7B**) at end-diastole, and in sub-valvular myocardium at end-systole (**Figure 7C**): As shown, MitraClip increased *E*_*rr*_ at end-diastole by 138 % in sub-valvular myocardium, and by 131% in baseline and LV volume augmented models respectively (both p<0.0001). MitraClip yielded even larger effects on endocardial end-diastolic *E*_*rr*_, as evidenced by increases of 230 % and 186 % in baseline and LV volume augmented models respectively (both p<0.001), corresponding to absolute increments of 0.052±0.035 and 0.044±0.049. Conversely, MitraClip decreased end-systolic *E*_*rr*_ at ES in lateral sub-valvular myocardium by 41.9 % and 45.1 % in baseline and LV volume augmented models respectively (both p<0.001) (**Figure 7C**). The effect of LV size was not significant at ED or at ES.

The quantitative effect of MitraClip on longitudinal strain (*E*_*ll*_) is shown in **Appendix Figure 1**. The pattern of effect on *E*_*ll*_ is similar to that of radial strain but inverted, meaning that longitudinal strain generally decreased post-procedure. Different than *E*_*rr*_, the effect of LV size was significant at ED and ES.

**Figure 6.**
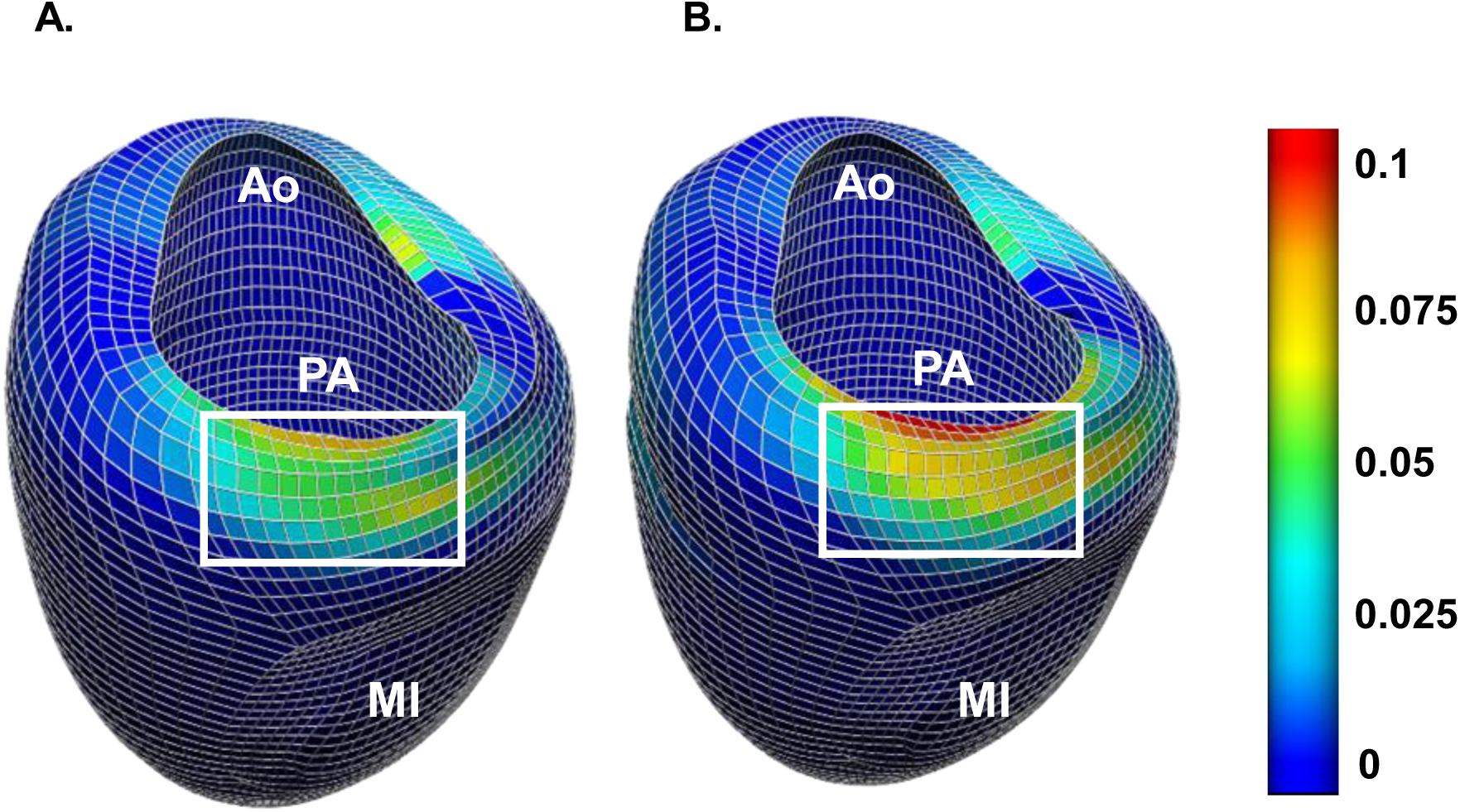
Color map of Δ in myocardial radial strain, *E*_*rr*_, at end-diastole after simulated complete grasp MitraClip in the baseline LV (**A**) and in a dilated LV (**B**). Ao – left ventricular outflow/ aorta region. PA – posterior mitral annulus. MI – myocardial infarction.

**Figure 7.**
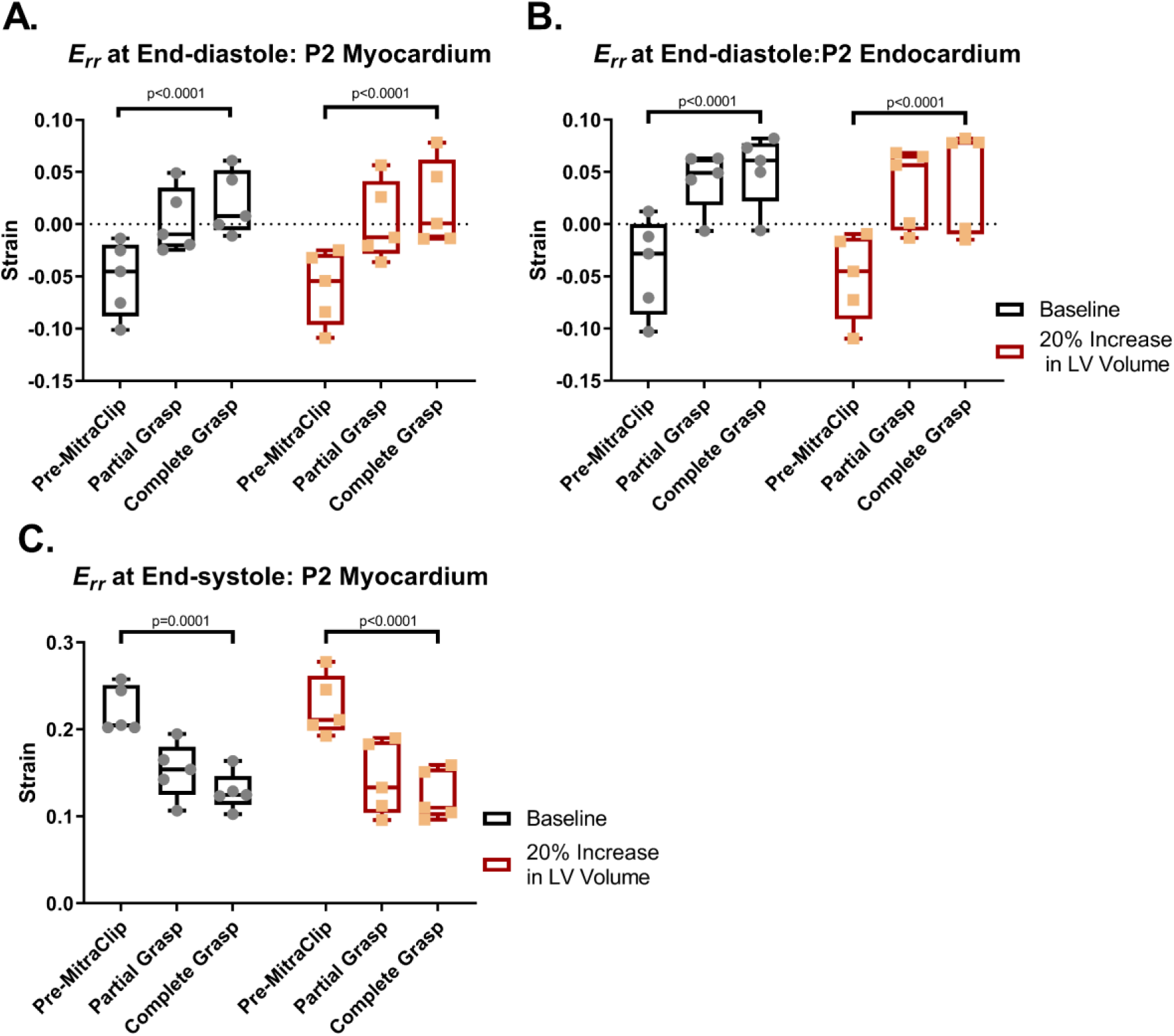
Radial strain, Err, after simulated partial and complete grasp MitraClip in the P2 sub-valvular myocardium at ED (A), P2 sub-valvular endocardium at ED (B) and P2 sub-valvular myocardium at ES (C) respectively. Err before and after MitraClip is relative to the pre-procedure unloaded state.

## Discussion

The principal finding of this study is that MitraClip implantation for FMR increases both end-diastolic and end-systolic mitral leaflet stress in the peri-device region: As a consequence, septo-lateral annular diameter was decreased and *E*_*rr*_ in the P2 sub-valvular myocardium was increased at end-diastole (ED) and decreased at end-systole (ES). Mechanical effects of MitraClip are augmented by in the enlarged LV, as leaflet stress and *E*_*rr*_ were further increased in ↑LV_VOLUME models at end-diastole.

### Leaflet stress

Regarding the underlying mechanism of MitraClip action, it should be noted that mitral leaflet stress is a function of leaflet and annular geometry, leaflet material properties, and loading conditions. MitraClip pulls the anterior and posterior leaflet edges together, thus inducing stress on the mitral leaflets that would be expected to increase in proportion to the amount of tissue gathered by the MitraClip (**Figure 8** illustrates proposed mechanism).

**Figure 8.**
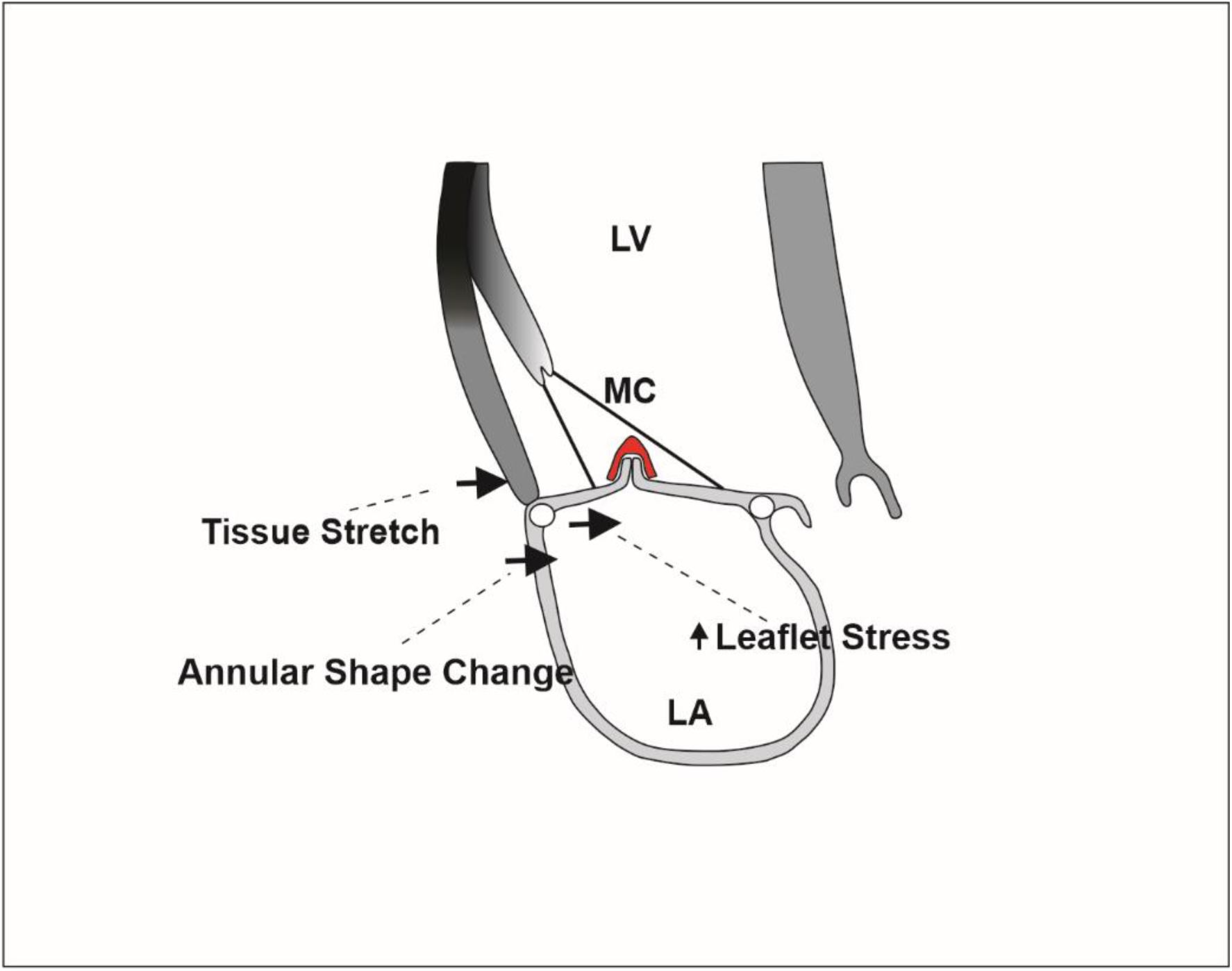
Diagram of the proposed mechanical effects of MitraClip application. The black arrows represent displacement of the posterior leaflet, lateral annulus and lateral sub-valvular LV wall caused by MitraClip application. The effects are to increase leaflet stress, decrease the septo-lateral annular dimension and stretch the sub-valvular myocardium. LV – left ventricle, LA – left atrium, MC – MitraClip

Increased LV volume is associated with annular enlargement and greater papillary muscle displacement. Simulation of LV volume effect in this study using ↑LV_VOLUME FE models confirms that pre-procedure increase in LV volume increases MitraClip-associated leaflet stress. Of note, the LV volume index at ES of sheep used in this study was approximately 40 ml/ m^2^ - a value lower than the volume index of most patients with FMR.

It is worth noting that increased trans-mitral gradient after MitraClip has been shown to predict poor composite outcomes including heart failure and death. [11] It is likely that those patients have leaflet stress and strain in the sub-valvular LV wall in excess of values determined in this study. Further studies are warranted to test these concepts.

### MitraClip durability

There is substantial evidence that mitral leaflet stress causes leaflet remodeling. Mitral leaflet thickening occurs with volume loading associated with pregnancy [28] and with leaflet tethering insufficient to cause MR [29] and also in sheep [30] and humans with FMR associated with increased LV sphericity, mitral tenting and annular size [31]. Along these lines, it is interesting that the EVEREST investigators found a difference between FMR and DMR in healing response when the MitraClip was explanted > 30 days after implantation. [32] Specifically, MitraClip implanted for FMR had thicker fibrous capsules and greater tissue bridge extending over the arms of the clip. [32]

There are also reports of acute leaflet perforation after MitraClip for FMR. [33] While usually considered a technical complication, acute perforation may be a function of leaflet stress since tissue rupture is more likely at high stress and strain levels. Taken together, these data support the concept that recurrent MR after MitraClip for FMR is impacted by device-induced leaflet stress.

### Procedure-related strain and post-MitraClip LV remodeling

Gathering of leaflet tissue by the MitraClip applies traction to leaflet, annulus and adjacent sub-valvular myocardium and subsequently increases leaflet stress, decreases the SL annular dimension and in turn increases radial strain, *E*_*rr*,_ in the adjacent sub-valvular myocardium as seen in **Figure 8.** The predominant effect on the LV wall is constraint of the myocardium which prevents wall thinning during diastole. As a consequence, systolic wall thickening is decreased, sarcomeres in the sub-valvular myocardium are not stretched and systolic wall thickening is decreased via the Starling mechanism.

It is known that volume and pressure overload associated stress and strain stimulate hypertrophy [34] and remodeling after myocardial infarction. [35] *In-vitro* studies suggest that a 10% strain level is sufficient for growth of myocyte sheets via serial sarcomere deposition. [36] MitraClip increases *E*_*rr*_ in the endocardium to near 5% at the end of diastole but it is unclear if this amount of diastolic thickening is sufficient to cause or accelerate LV remodeling in the sub-valvular myocardium.

Pantoja et al. studied the effect of UA on procedure related strain in the LV wall and showed that UA significantly increased longitudinal strain in the proximal lateral LV wall. [16] Specifically, the largest effect of UA on *E*_*ll*_ occurred in the proximal-lateral endocardial surface, where strains increased from 0.0345 ± 0.0206 to 0.1117 ± 0.0480 (p=0.0057). Pantoja et al., did not measure the effect of UA on annular shape but when compared with MitraClip, the magnitude of annular reduction was likely significantly larger and annular reduction extended across the entire posterior annulus.

### Amount of MitraClip tissue grasp

MitraClip NTR with both partial and complete grasp was simulated in this study where compared to the partial grasp, the complete grasp had overall larger effect on leaflet stress, SL dimension and *E*_*rr*_ in LV sub-valvular myocardium. Based on this observation, we anticipate that compared to MitraClip NTR (9 × 5mm), MitraClip XTR (12 × 5mm) will cause even larger leaflet stress concentration and more positive *E*_*rr*_ at ED and cause an even larger reduction in the SL dimension at ED.

### Study limitations

The primary limitation of this study is that our calculations were not optimized with experimental data obtained after MitraClip. On the other hand, SL annular dimension at end-diastole decreases between 2.8 to 4 mm in patients with FMR after MitraClip [37-39] and Lurz et al. found that the MRI-measured radial strain averaged across all sectors at the mitral valve level decreased from 15.4 to 9.6% after MitraClip in a cohort of mostly FMR patients. [40] These findings show that our simulations are in agreement with clinical results.

Second, the MR jet location was not measured in the sheep used in the current study. [18] Three-dimensional echocardiography has been performed on sheep with postero-lateral MI [41, 42] but MR jet location has not been reported. [41, 42] On the other hand, Hidalgo et al found that 41% in patients with FMR undergoing MitraClip had an MR jet in the A2P2 region and 45.5% had a jet that spanned the A2P2-A3P3 region. [38] This supports the contention that postero-lateral MI is associated with leaflet restriction along the entire A2P2 A3P3 section rather than at A2P2 or A3P3 separately. Location of the virtual MitraClip at A2P2 in the current study is therefore reasonable.

Finally, leaflet stresses at ED and ES were examined in the current study, although Rabbah et al. observed that leaflet stress was greatest during peak systole. [43] Future planned analyses will include stress at peak systole.

## Conclusions and Future Directions

Finite element simulations performed in this study demonstrate that MitraClip increases peri-MitraClip leaflet stress at both ED and ES when performed for FMR. Subsequent traction on the mitral annulus causes the SL dimension to decrease and endocardial strain in the adjacent LV wall at ED is increased. Further, mechanical effects of MitraClip on leaflet stress and sub-valvular LV strain are augmented in the enlarged LV Increased mitral leaflet stress and procedure-related strain in the sub-valvular myocardium may contribute to mitral leaflet and LV remodeling after MitraClip. Future clinical studies are warranted to clinically test device-induced mechanical effects as predictors of procedural success for patients undergoing MitraClip.

## Acknowledgements

This study was supported by NIH grants R01-HL128099/Levine, R01 HL141917/Levine, R01-HL63348/Ratcliffe and R01-HL128278/Weinsaft.

This support is appreciated.

---

**Appendix Figure 1.**
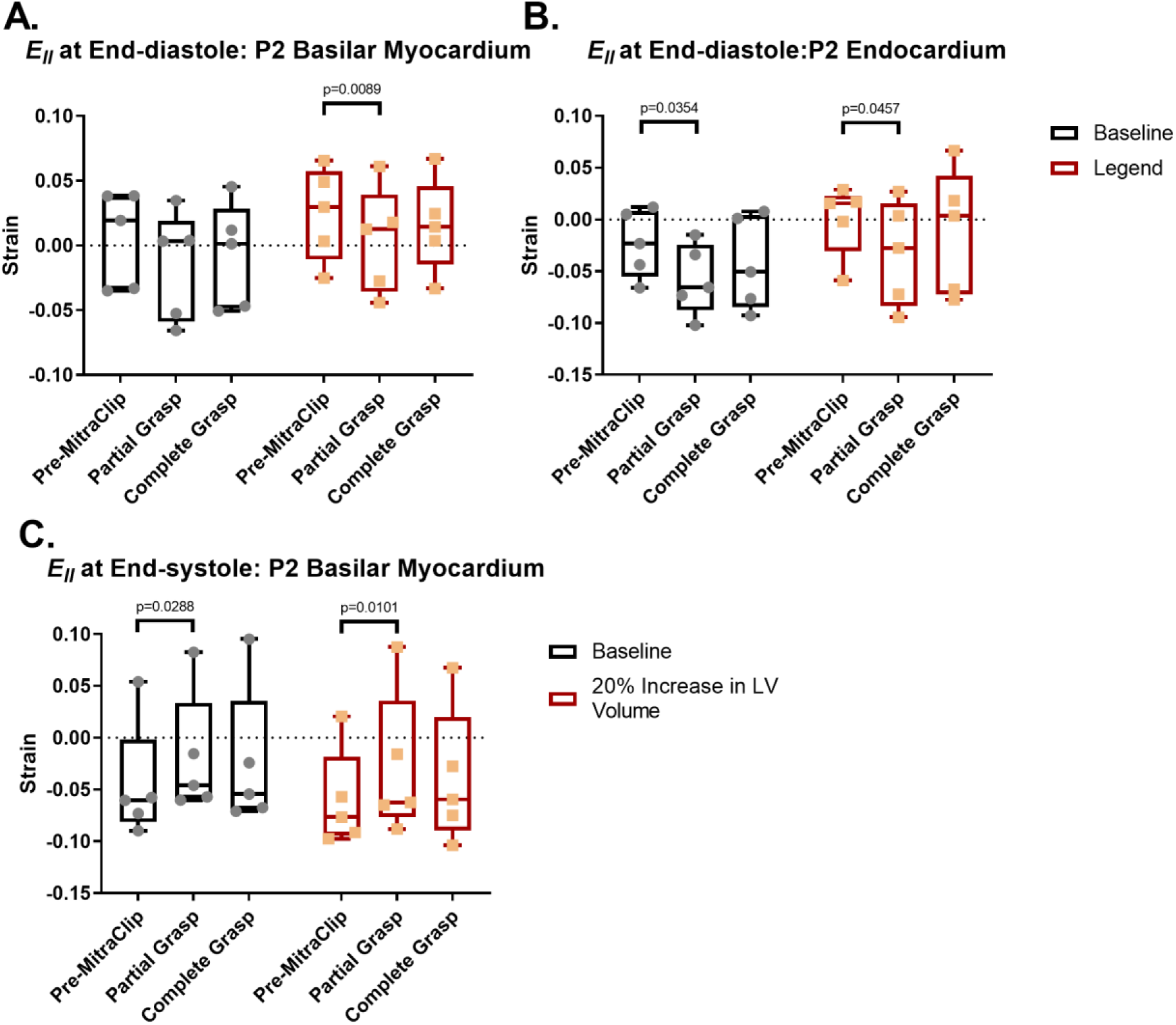
Longitudinal strain, Ell, after simulated partial and complete grasp MitraClip in the P2 sub-valvular myocardium at ED (A), P2 sub-valvular endocardium at ED (B) and P2 sub-valvular myocardium ES (C) respectively. Ell before and after MitraClip is relative to the pre-procedure unloaded state.

